# CrispRVariants: precisely charting the mutation spectrum in genome engineering experiments

**DOI:** 10.1101/034140

**Authors:** Helen Lindsay, Alexa Burger, Berthin Biyong, Anastasia Felker, Christopher Hess, Elena Chiavacci, Jonas Zaugg, Carolin Anders, Martin Jinek, Christian Mosimann, Mark D. Robinson

## Abstract

CRISPR-Cas9 and related technologies efficiently alter genomic DNA at targeted positions and have far-reaching implications for functional screening and therapeutic gene editing. Understanding and unlocking this potential requires accurate evaluation of editing efficiency. We show that methodological decisions for analyzing sequencing data can significantly affect mutagenesis efficiency estimates and we provide a comprehensive R-based toolkit, CrispRVariants and accompanying web tool CrispRVariantsLite, that resolves and localizes individual mutant alleles with respect to the endonuclease cut site. CrispRVariants-enabled analyses of newly generated and existing genome editing datasets underscore how careful consideration of the full variant spectrum gives insight toward effective guide and amplicon design as well as the mutagenic process.

---

Genome engineering technologies are developing at a rapid pace. The most prominent methods based on bacterial CRISPR (clustered regularly interspaced short palindromic repeats) systems couple a predesigned 20bp short guide RNA (e.g. sgRNA) with a protein (e.g., Cas9^1^, Cpf1^2^) carrying nuclease activity. The sgRNA targets the nuclease machinery to the genomic locus of choice, resulting in double-stranded breaks in DNA. Typically, a number of bases are inserted or deleted in a stochastic manner as the two DNA ends are rejoined by non-homologous end joining (NHEJ)^3^. Optionally, donor DNA can be introduced and integrated between the breakpoints^4,5^. The result is an “edited” genome sequence at a chosen location. The growing application of CRISPR systems and related technologies for genome editing are driven by their seemingly universal high efficiency in an increasing number of cell types and animal systems^5–7^ as well as the significant potential to establish causal links between genotype and phenotype.

In *vivo* CRISPR applications, where multiple cells undergo independent rounds of mutagenesis and local NHEJ, generate particularly heterogenous sequencing data sets. Existing tools for the analysis of mutagenesis sequencing data report aggregated variant summaries (**CRISPR-GA**^8^, **CRISPResso**^9^) and are unsuited for applications that consider the entire, complex mutation spectrum, e.g. quantifying mosaicism^10^ and allele-specific genome editing^11^. To facilitate such analyses, we have developed **CrispRVariants**, an R-based toolkit for quantifying and visualising individual variant alleles from either traditional Sanger sequencing or high-throughput CRISPR-Cas9 mutagenesis sequencing experiments. **CrispRVariants** can be easily used to create a variant allele summary plot (Figure 1) and accompanying table of counts. Individual variants can be removed, allowing allele-specific analysis and adjustment for heterozygosity. By localising variant alleles with respect to the nuclease cut site instead of the PCR amplicon, **CrispRVariants** enables immediate comparison of variant spectra between target locations (Supplementary Note 1). This level of resolution enables users to directly relate variant genotypes to observed phenotypes and predict downstream effects of variants, such as protein structural changes or loss or gain of transcription factor binding sites when targeting non-coding sites (see examples in the **CrispRVariants** User Guide and Reference Manual). Figure 1 summarizes several zebrafish embryos injected with an sgRNA targeting *ENSDARG00000079624 (wtx)*, which results in a variety of alleles, some of which reoccur independently in multiple embryos. Importantly, visualization of variant alleles facilitates the detection of sequencing or alignment errors and previously-unknown genetic variation. We designed **CrispRVariants** with interactivity in mind, explicitly allowing users to detect problems and filter sequences appropriately before estimating mutation efficiency. The accompanying web tool, **CrispRVariants**Lite, which is suitable for smaller-scale experiments, can be accessed via the website or downloaded and run locally, allowing users without bioinformatics expertise to examine and plot their data.

**Figure 1.** The **CrispRVariants** `plotVariants` function summarizes variant types, locations and frequency across multiple clones from several injected animals. This function returns a **ggplot2**-based allele summary plot consisting of (1) a schematic of the target site location relative to the neighboring transcripts, (2) an alignment of the consensus sequence for each variant combination to the reference sequence, and (3) a heat map showing the frequency of the variants across samples (the heatmap can be plotted also with frequencies). Inserted sequences are shown below the alignments, with large insertions indicated by the corresponding symbol. In this example, columns in the heat map represent sequences cloned from different embryos, with column labels colored by the embryonic phenotype (black = uninjected, blue = wild-type-like, orange = developmental abnormalities or “monsters”, green = heart phenotype).

Distinguishing low-frequency mutation events from sequencing errors is challenging and while most amplicon sequencing studies lack ground truth, sequencing the offspring of mutagenized animals results in a situation where two different alleles are expected. In Supplementary Note 2, we show examples of sequencing errors and alignment uncertainty that affect the size, placement and ultimately variant classification (i.e., whether in-frame or not) of two germline mutant cohorts. Sequencing errors and genetic variation confound mutation efficiency estimation; for example, sequence polymorphisms in the targeted locus affect sgRNA binding and may lead to underestimation of the true editing efficiency. In Supplementary Note 3, we highlight unappreciated genetic variation in a recent study^12^ as well as off-target sgRNAs that lack a canonical PAM sequence. We show through simulation that **CrispRVariants** matches or outperforms existing tools in estimating mutation efficiency (Supplementary Note 4). Notably, blind data processing decisions contribute substantially to the differences between **CrispRVariants** and other available tools. We include with **CrispRVariants** a small synthetic benchmarking data set containing several types of commonly observed variants to facilitate transparent data processing.

Despite overwhelming evidence that data preprocessing choices affect variant calling in exome and whole-genome sequencing studies^13^–^15^, their role in estimating mutagenesis efficiency has been largely neglected. Amplicon sequencing data may be aligned locally to the expected amplicon sequence (e.g. Gagnon et al.^5^, **CRISPR-GA**^8^, **CRISPResso**^9^), in which case pooled reads must first be separated, or globally to an entire reference genome (**AmpliconDIVider**^16^, **CrispRVariants**). Strategies that combine local and global alignment (**CRISPResso** (**Pooled**)^9^) or avoid separating reads by aligning to the set of all amplicons^17^ are also possible. Inappropriate alignment and preprocessing settings can have a significant impact on allele counts and efficiency estimates. In the most extreme case, tandem repeats and homology within an amplicon resulted in efficiency estimates that differed by 91% between methods (Supplementary Note 5). Local alignment strategies are vulnerable to mis-counting off-target reads. For example, **BLAT**^18^ local alignment (used in **CRISPR-GA**) can result in efficiency estimates that differ by more than 30% from estimates from global alignments (Supplementary Note 5). Stringency criteria when merging paired-end reads or dividing reads by PCR primers (as done in Shah et al.^12^) can further affect mutation efficiency estimates (Supplementary Notes 6 and 7). Specifically, altering the percentage overlap required for merging from 100% (as in Shah et al.^12^) to 90% changed the efficiency estimate for one guide by 65%. **CrispRVariants** separates data preprocessing from mutation quantification, allowing critical parameters to be carefully selected and tailored to the experimental design (see Methods). By aggregating variant alleles instead of looking at and interpreting the full observed spectrum, existing tools make it difficult to assess whether appropriate bioinformatic decisions (alignment, merging, separation of reads) have been made; **CrispRVariants** facilitates this visual, interactive and iterative process.

In summary, the **CrispRVariants** package offers precise, transparent and reproducible preprocessing of low- and high-throughput amplicon sequencing experiments, providing easy visualizations of variant alleles across samples and allows careful calculation of the efficiency, given all the complexities and confounders. The resulting allele summary plots (Figure 1) provide a compact representation of deletions as well as potentially complex insertions. The **CrispRVariants** package is extensible, fully customizable and facilitates interactive and iterative analyses. At present, **CrispRVariants** is targeted toward CRISPR-Cas9 systems, where alignments are placed in context of the PAM sequence with the expected cut site highlighted; however, the framework can readily be applied to other mutagenesis systems. The software is available from **Biocon-ductor**^19^ (https://www.bioconductor.org/packages/CrispRVariants) and interfaces seamlessly with existing **Bioconductor** infrastructure.

## Methods

**CrispRVariantsLite** is available online via: http://imlspenticton.uzh.ch:3838/CrispRVariant the code and instructions for local installations is available from: https://github.com/markrobinsonuzh/CrispRVariantsLite/.

For both low- or high-throughput sequencing analysis, the main entry point to **CrispRVariants** is a set of sequences aligned to a reference genome in BAM (binary alignment) format. Reads that cannot be represented as a single linear alignment are instead represented by some alignment tools as multiple “chimeric” alignments. We find that some chimeric reads are genuine variants (Supplementary Note 8) and recommend the use of a chimera-aware aligner. In current pipelines, we use **BWA MEM**^20^ with default parameters. The choice of aligner can substantially affect the mutation efficiency estimates (Supplementary Note 5). Applied Biosystems Sanger sequencing data, commonly available in AB1 file format, can be easily converted to FASTQ format for mapping; **CrispRVariants** uses the **sangerseqR**^21^ package to perform this conversion. The entry points for **CrispRVariantsLite** include a ZIP file of BAM files (sets of already mapped reads), a ZIP file of directories with AB1 files or a ZIP file of FASTQ files (file size restrictions apply).

**CrispRVariants** can work directly with pooled amplicon sequencing data. Reads are assigned to the correct amplicon either by an alignment spanning the amplicon region almost exactly (strict), or by any base mapped to the unique portion of an amplicon (relaxed). Because of high error frequency, the endpoints of Illumina MiSeq data are often clipped by aligners. We extrapolate the mapped region to include clipped regions when matching amplicons. This dividing strategy is suitable for paired-end reads where both reads span the entire amplicon, or for merged paired-end reads. In cases where unique mapping to a single amplicon is insufficient to assign reads, alignments may be filtered in **R** and passed directly to **CrispRVariants** as a GenomicAlignments^22^ object. **CrispRVariants** can collapse paired reads by checking for concordant variants in the vicinity of the cut site. However, if merging criteria are not overly strict, we find that merging reads prior to mapping improves speed without affecting efficiency estimates (Supplementary Note 6). Chimeric reads are assigned to all overlapping amplicons, however, to be counted as a variant the mapped endpoint of one aligned segment must be close to the specified cut site. This criterion excludes PCR artifacts, such as primer dimers. Chimeric read sets are grouped into an “Other” category. For amplicons with a non-trivial fraction of “Other” reads, additional exploratory analyses are available within the software (see software vignette).

Once assigned, read alignments are narrowed to the target region (i.e., the user-specified local genomic region around the guide’s target site). Reads that do not span the target region are discarded and reads that match the reference sequence are recorded as “no variant”. Insertions and deletions (indels) are then localized in a strand-aware manner, labeled and counted; a 3 base pair deletion starting 2 bases upstream of the target location is designated “-2:3D”. Downstream variants are numbered similarly by their leftmost base. Reads that do not contain an indel can additionally be separated by the presence of single nucleotide variants (SNVs). By default, the zero point is at base 17 of a 23 bp sgRNA, i.e. the endonuclease cut site. The user is free to specify: i) the target region; ii) the corresponding zero point; and iii) a window within the target region for calling SNVs.

**Supplementary analyses** For the Supplementary analyses, we use data from Shah et al.^12^, Burger et al.^23^ and Cho et al.^24^. The performance of **CrispRVariants** (version 0.9.2), **CRISPResso** (version 0.8.2), **CRISPR-GA** and **AmpliconDIVider** was compared under a range of scenarios. Where not otherwise specified, the data is from Shah et al.; **CrispRVariants** and **AmpliconDI Vider** were run after **BWA MEM** alignment and **CRISPResso** was run in single amplicon mode. Further information about the data used is in Supplementary Note 9.

## Acknowledgements

HL and MDR acknowledge funding from the European Commission through the 7th Framework Collaborative Project RADIANT (Grant Agreement Number:305626).

Competing Interests The authors declare that they have no competing financial interests.

